## Supplement for "Long-read sequencing transcriptome quantification with lr-kallisto"

### Supplementary Figures
### Long-read sequencing transcriptome quantification with lr-kallisto

**Rebekah K. Loving[1], Delaney K. Sullivan[1,2], Fairlie Reese[3,4], Elisabeth Rebboah[3,4], Jasmine Sakr[3,4], Narges Rezaie[3,4], Heidi Y. Liang[3,4], Ghassan Filimban[3,4], Shimako Kawauchi[3], Conrad Oakes[1], Diane Trout[1], Brian A. Williams[1], Grant MacGregor[3], Barbara J. Wold[1], Ali Mortazavi[3,4], and Lior Pachter[1,5]**

[1]Division of Biology and Biological Engineering, California Institute of Technology, Pasadena, CA 91125
[2]UCLA-Caltech Medical Scientist Training Program, David Geffen School of Medicine, University of California, Los Angeles, Los Angeles, CA, 90095, USA
[3]Developmental and Cell Biology, University of California Irvine, Irvine, USA
[4]Center for Complex Biological Systems, University of California Irvine, Irvine, USA
[5]Department of Computing and Mathematical Sciences, California Institute of Technology, Pasadena, CA 91125

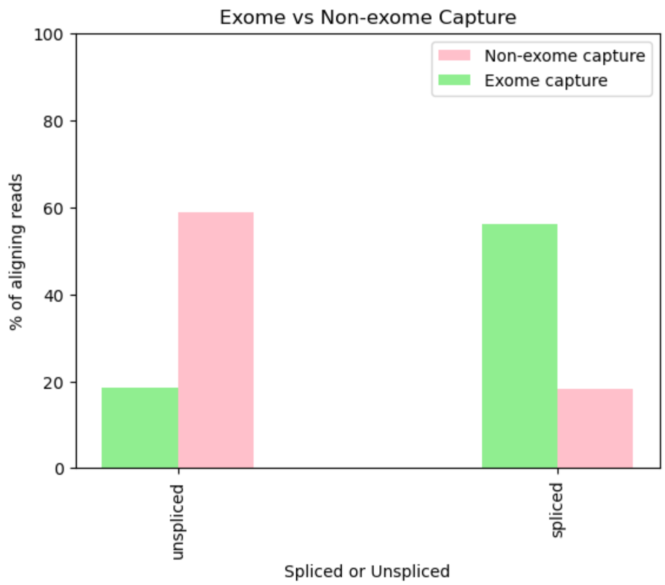


Supplementary Fig. 1a: Comparison of the percent of reads mapping as spliced vs. unspliced reads with and without exome capture.

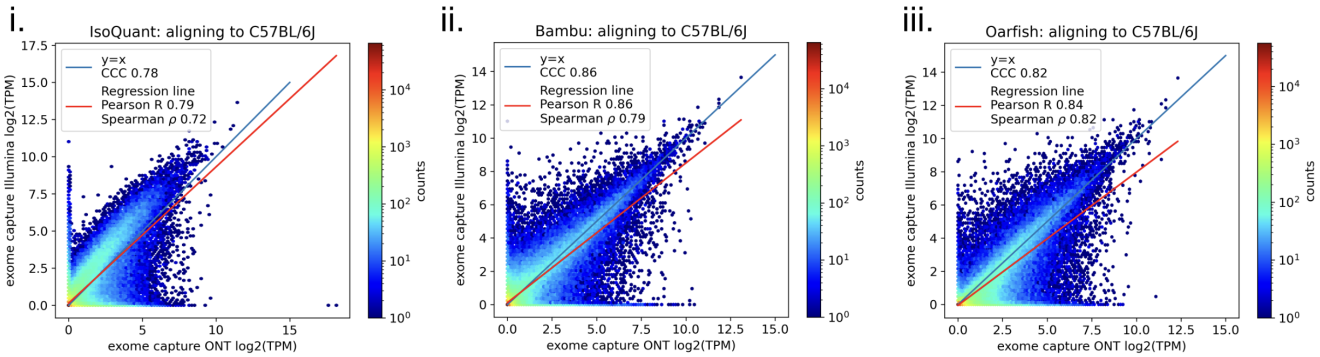


Supplementary Fig. 1b: Quantifications of the C57BL/6J exome capture samples with Bambu, IsoQuant, and Oarfish.

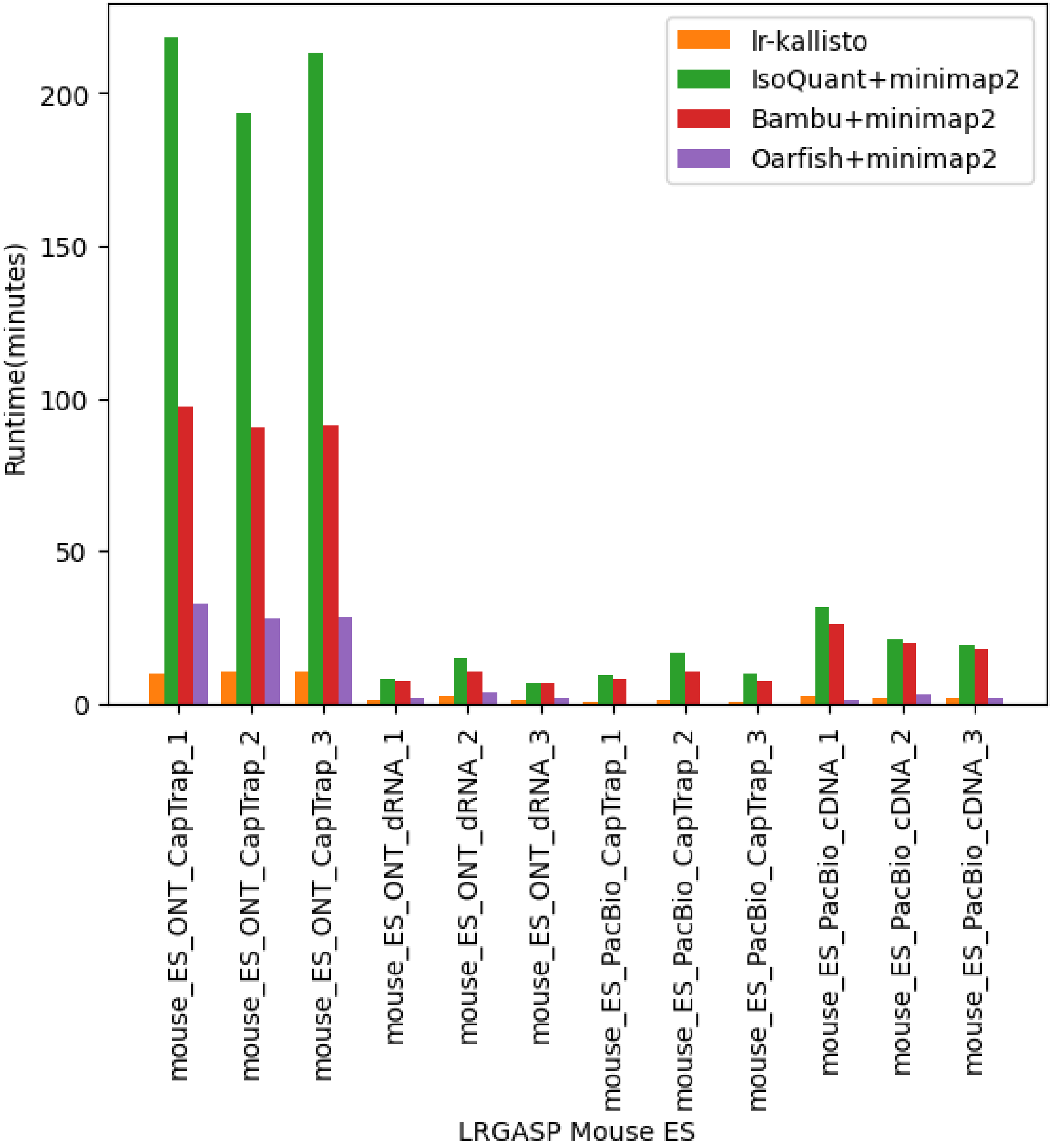


Supplementary Fig. 1c: Runtime performance comparisons for lr-kallisto, IsoQuant, Bambu and Oarfish.

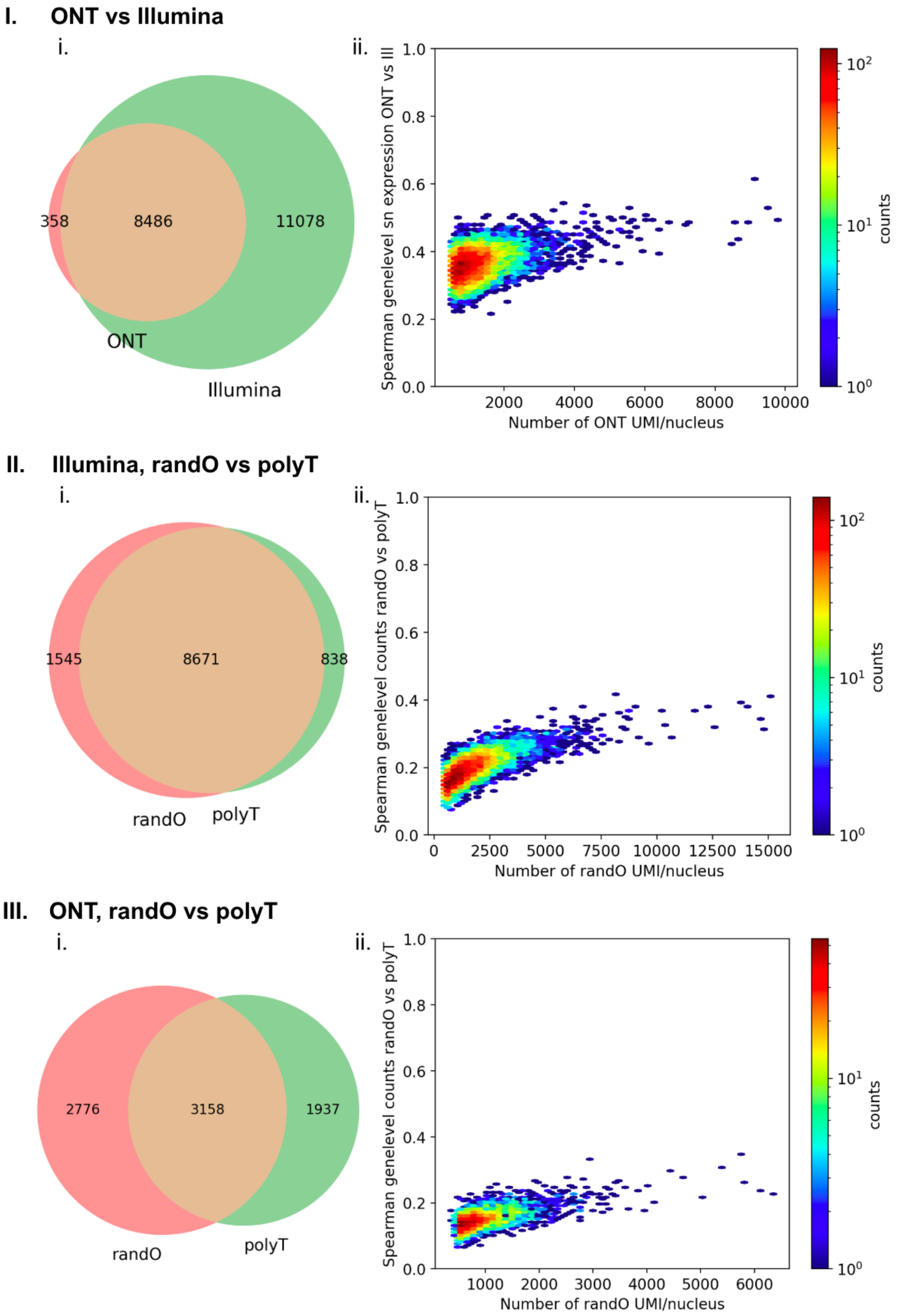


Supplementary Fig. 1d: I. i. Venn diagram of barcodes in ONT and Illumina. ii. Number of ONT UMI/nucleus vs Spearman correlation between ONT and Illumina single-nucleus gene level counts. II. i. Venn diagram of barcodes in Illumina random oligo (randO) and Illumina poly dT. ii. Number of randO UMI/nucleus vs Spearman correlation between Illumina randO and Illumina polydT single-nucleus gene level counts. III. i. Venn diagram of barcodes in ONT random oligo (randO) and ONT poly dT. ii. Number of randO UMI/nucleus vs Spearman correlation between ONT randO and ONT polydT single-nucleus gene level counts.

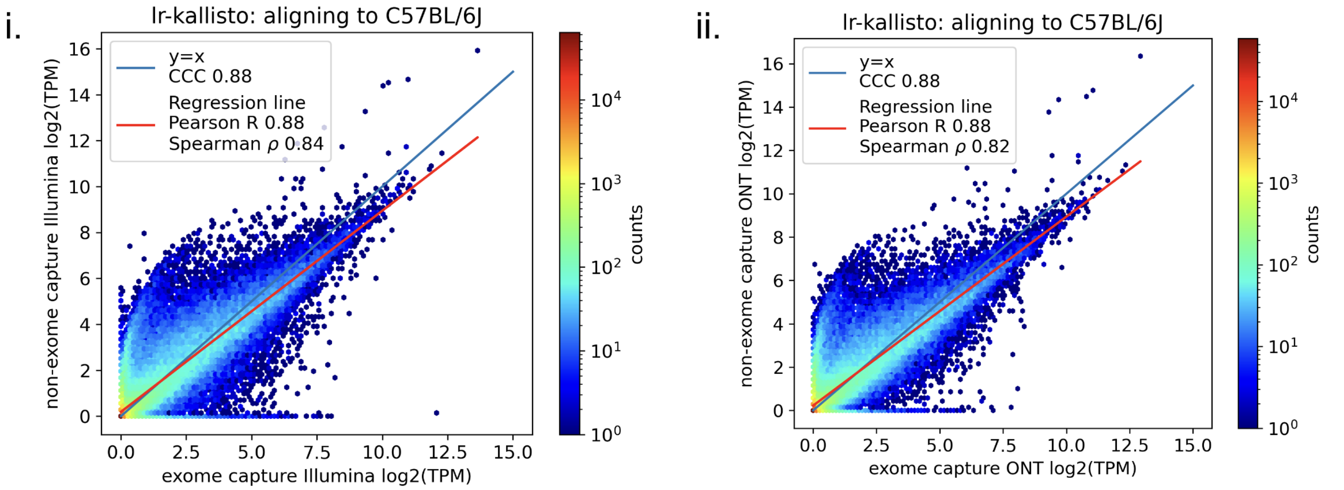


Supplementary Fig. 1e: Contrast of non-exome vs exome capture in Illumina and ONT.

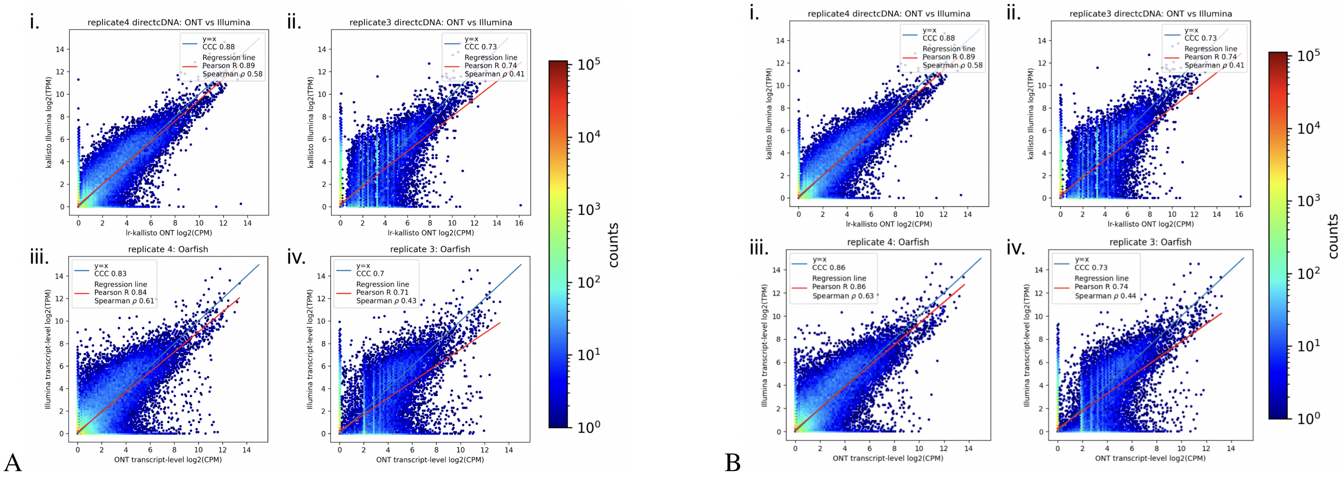


Supplementary Fig. 2a: Performance of lr-kallisto on ONT sequenced direct cDNA libraries from the HCT116 cell line, where panel A is with Oarfish v0.3.1 and panel B is with Oarfish v0.5.1.

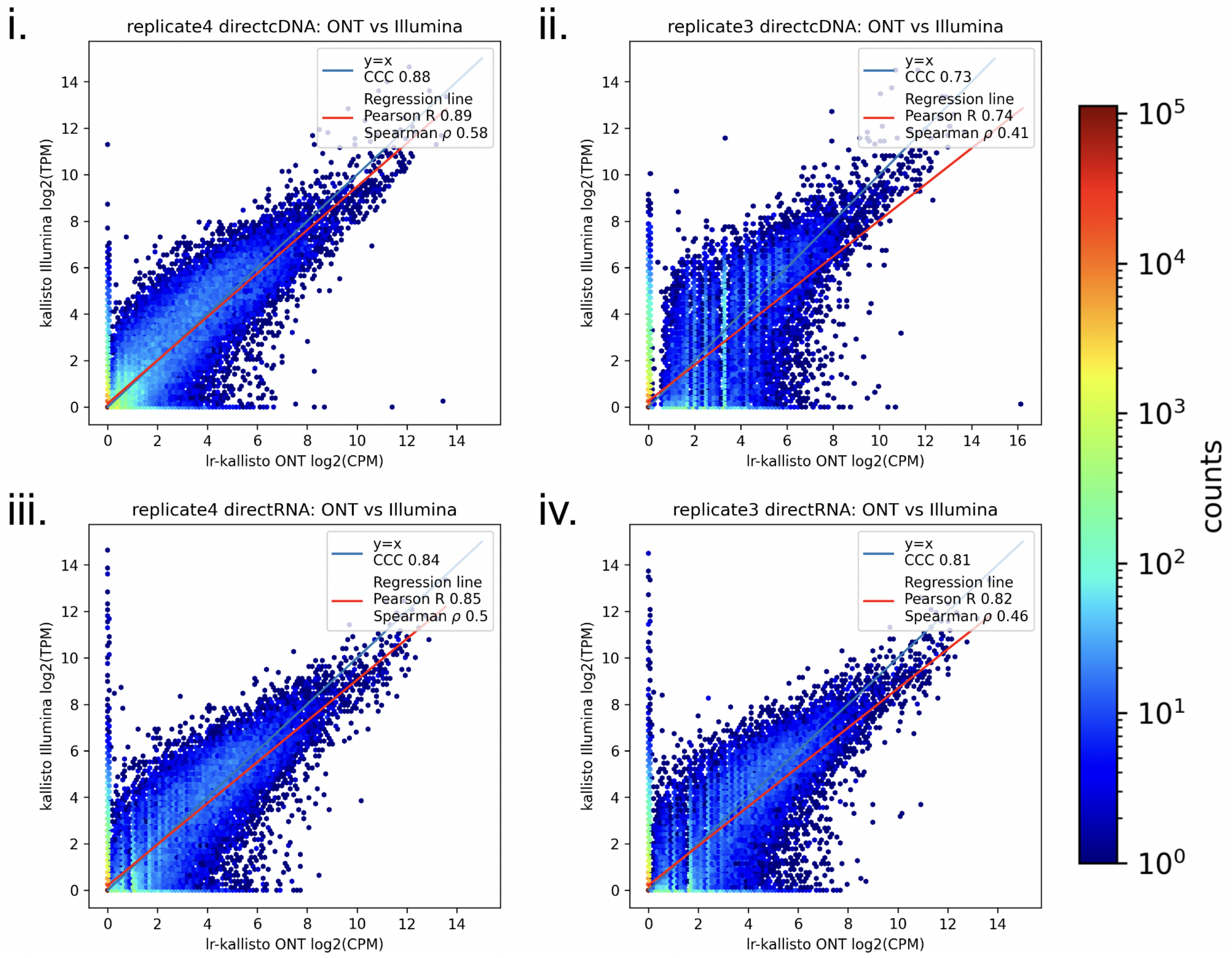


Supplementary Fig. 2b: Comparison of lr-kallisto on ONT sequenced HCT116 cell line libraries generated with directRNA and direct cDNA between two replicates in each.

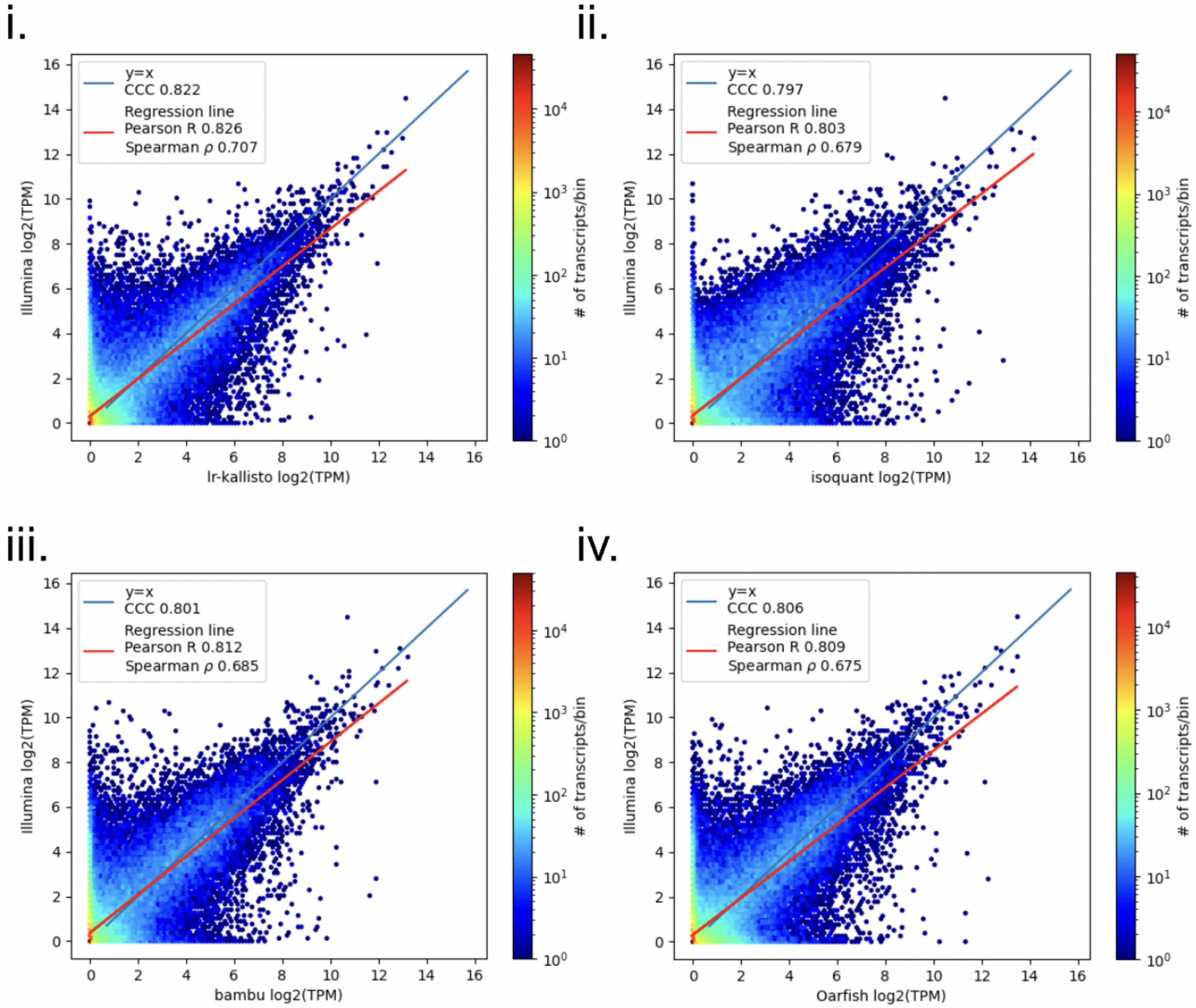


Supplementary Fig. 2c: Evaluation of Bambu, IsoQuant, lr-kallisto, and Oarfish on mouse cortex high-depth PacBio data.

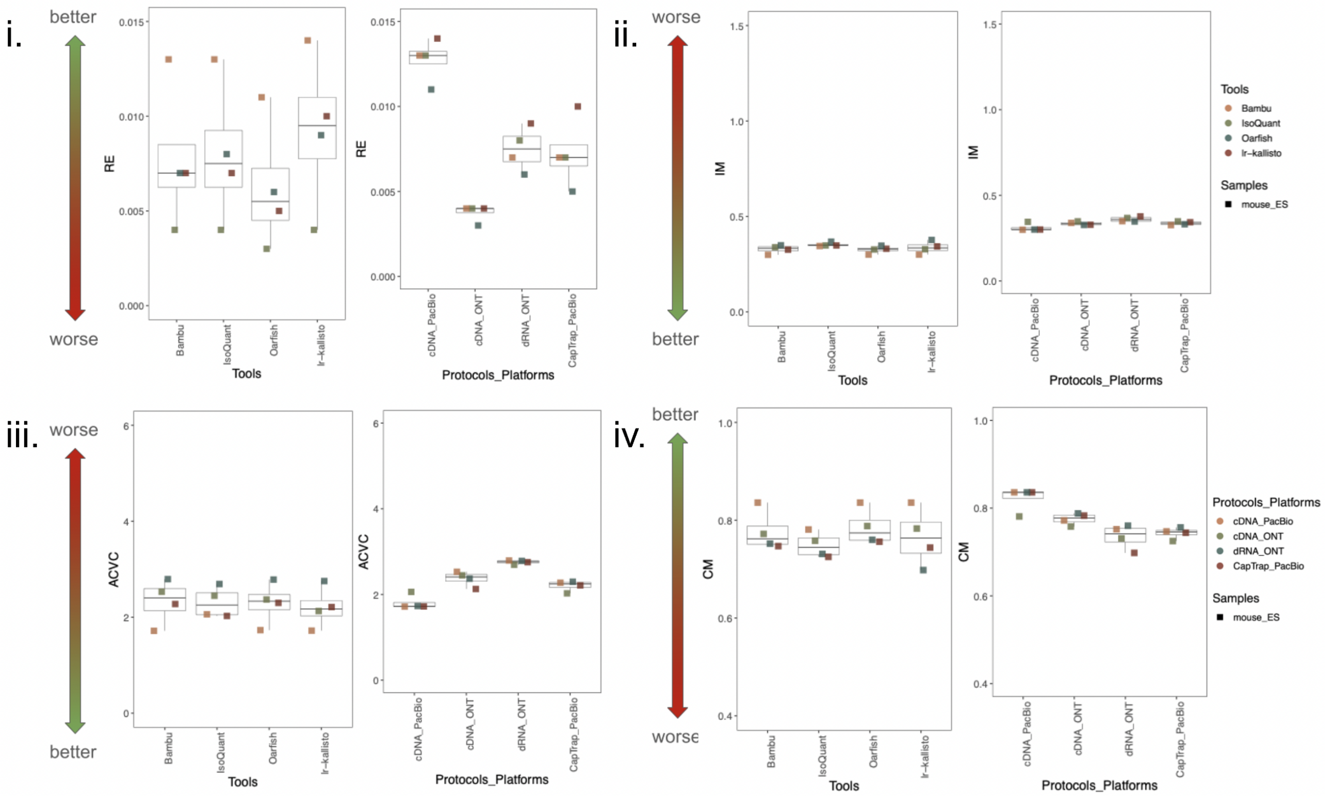


Supplementary Fig. 3: Evaluation of lr-kallisto, Bambu, IsoQuant and Oarfish according to LRGASP challenge 2 metrics in Mouse ES cells.

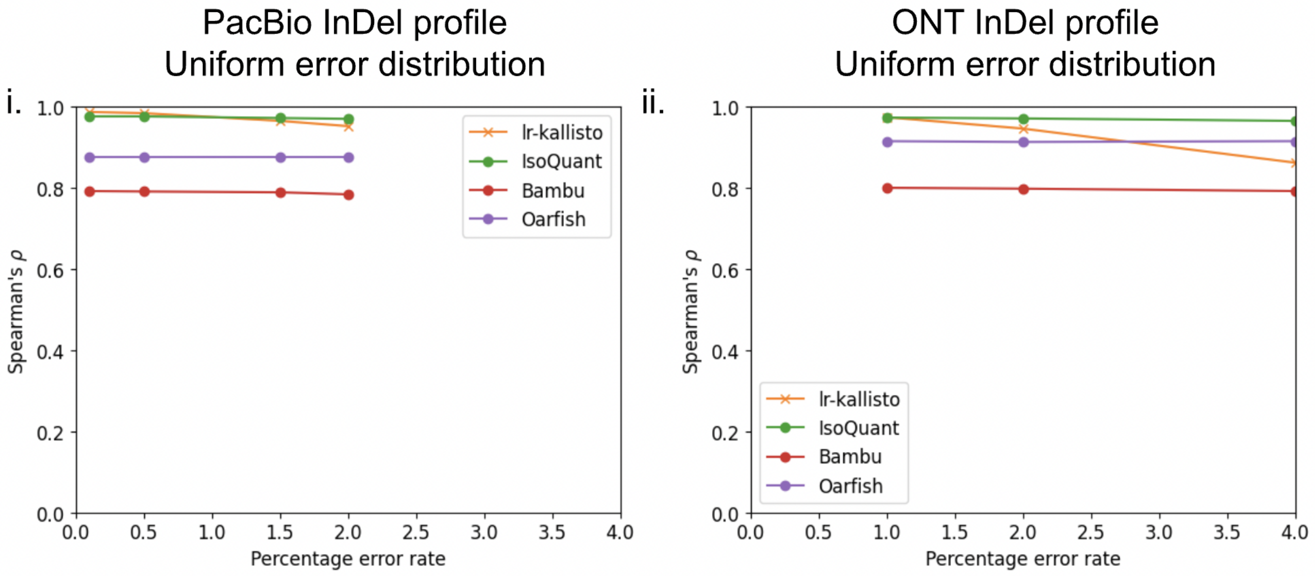


Supplementary Fig. 4a: Benchmarks of Bambu, IsoQuant, lr-kallisto, and Oarfish on simulations with a range of error parameters.

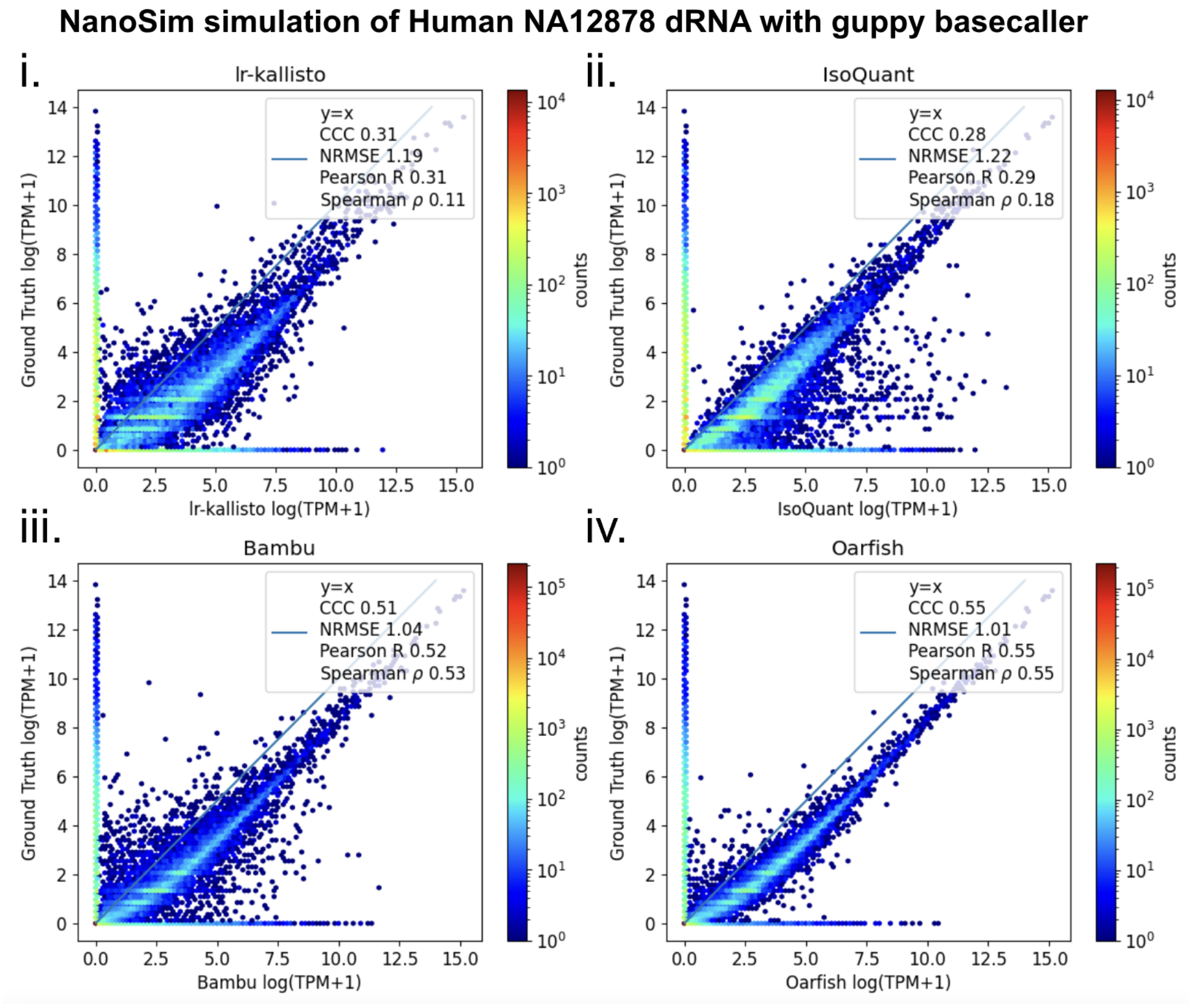


Supplementary Fig. 4b: Performance on all annotated transcripts at ONT 11.2% sequencing error rate.

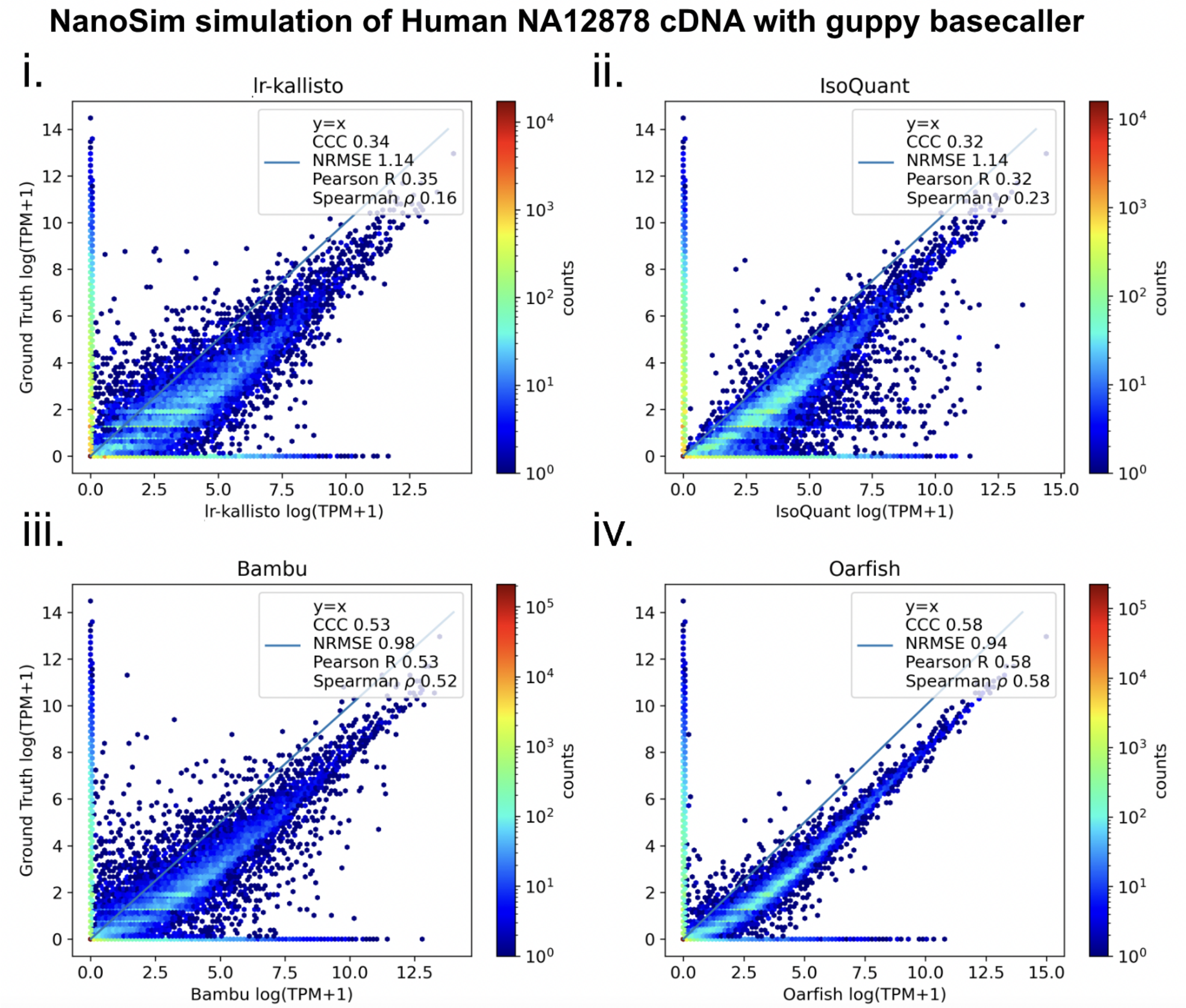


Supplementary Fig. 4c: Performance on all annotated transcripts at ONT 15.2% sequencing error rate.

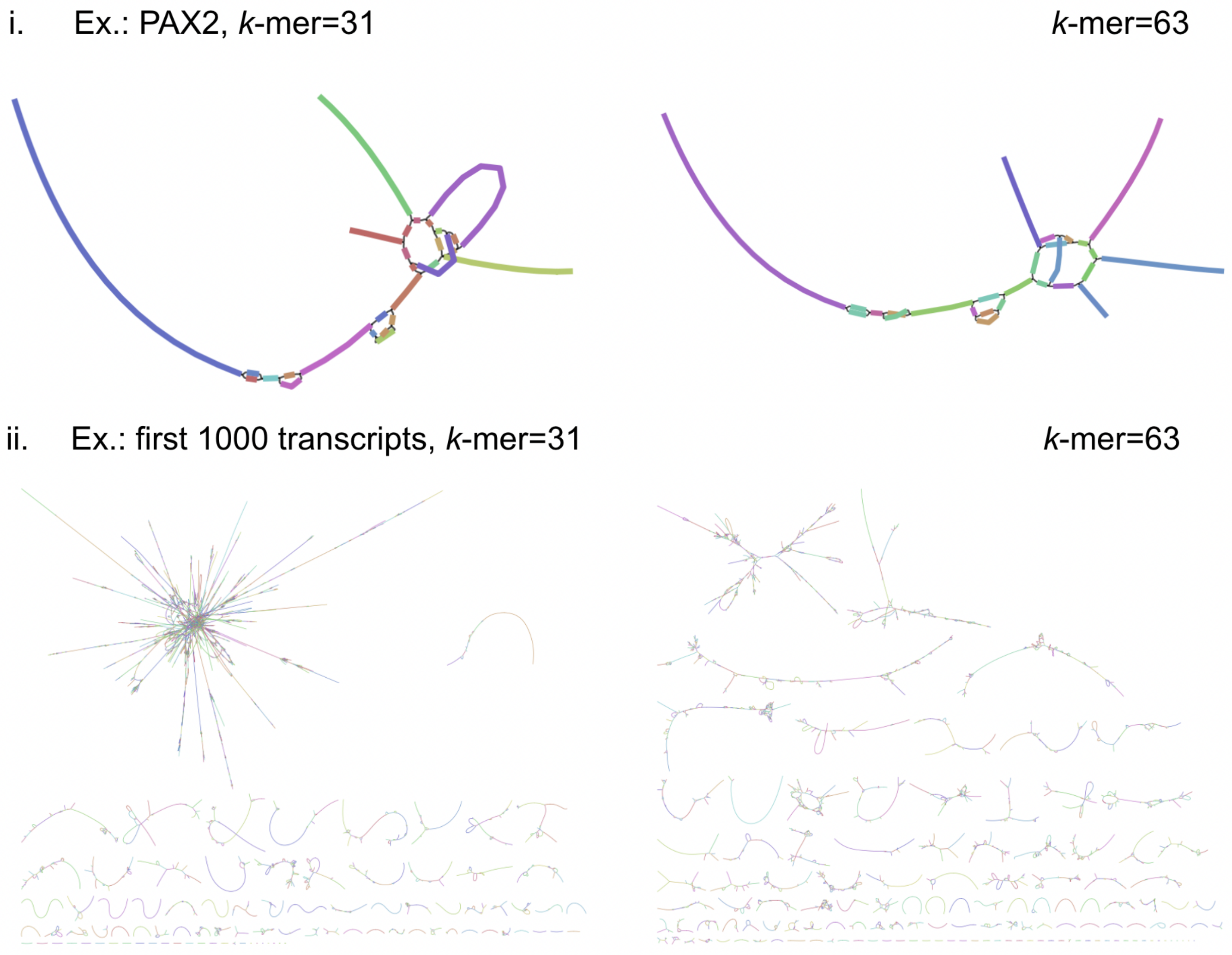


Supplementary Fig. 5: lr-kallisto transcript de Bruijn Graph bandage plots.
